# Hyper-mutational processes provide a head-start for weak cancer drivers: explaining atypical *KRAS* variants

**DOI:** 10.1101/2025.01.29.635499

**Authors:** Michael D. Nicholson, Ian Tomlinson

## Abstract

Hypermutant cancers frequently contain atypical driver gene variants, which may provide sub-optimal oncogenic advantages. The reason why weak cancer drivers prevail and are not outcompeted by stronger alternatives is unclear. Here, using mathematical modelling, we show that aberrant mutational processes alone can account for the detection of weak, rather than canonical, driver mutations. We find that simply increasing the mutation rate per cell division, without altering selection or mutational biases, can lead to the dominance of weak drivers due to rapidly occurring further driver events. This effect can be further enhanced by mutational biases towards specific nucleotide sequence contexts. Focusing on *POLE*-mutant (DNA polymerase epsilon proofreading-deficient) colorectal cancers, we quantify the mutation bias for varied *KRAS* drivers under both *POLE-*mutant and non-hypermutant mutational processes. In *POLE*-mutant cancers, the combination of the bias coupled with an elevated mutation rate is sufficient to explain the enrichment of atypical *KRAS* drivers. Furthermore, model predictions are consistent with the observed prevalence of atypical *KRAS* drivers observed in mismatch repair deficient colorectal cancer. Thus, differential selection across these cancer types need not be invoked to explain the variation in driver mutations. Our study clarifies the interplay of mutation and selection during the evolutionary dynamics of tumourigenesis.

**Author Summary:** Considerable variability exists in the effectiveness of cancer driving mutations to promote cancer. Mutations which are thought to be less effective, ‘weak’ driver mutations, are commonly observed in DNA repair deficient, ‘hypermutant’ cancers, for reasons that remain unclear. Through mathematical modelling, we detail conditions such that accelerated mutation rates, combined with the biassing of mutations towards certain DNA locations, leads to the dominance of weak driver mutations within a tumour. In contrast to classical mathematical models of cancer initiation, we find that heightened mutation burden alone can alter which driver mutations are to be expected. Applying our model to the canonical oncogene *KRAS*, we show that the occurrence of weak driver mutations is well explained by the aberrant molecular processes of hypermutant colorectal cancers. Differential effectiveness of the ‘weak’ drivers in hypermutant cancers is not required. Our study clarifies how mutation and selection interact during the evolutionary dynamics of tumourigenesis.

## Introduction

An overarching goal of cancer genomics is to understand why certain driver mutations are observed in specific cancers (Vogelstein et al. 2013) Given the varied micro-environments and epigenetic landscapes of tumours, distinct drivers are perhaps to be expected between cancers in different tissues (Poulin et al. 2019; Haigis et al. 2019). Less expected is that cancer subtypes within a given tissue - where subtypes are delineated by measures as simple as mutation burden - have divergent driver variants (Muzny et al. 2012; Cornish et al. 2024; Touat et al. 2020; Kim et al. 2022; Tan et al. 2022; Levine, D., The Cancer Genome Atlas Research Network 2013). For example, in the ‘hypermutant’ class of colorectal cancers (CRCs), the most frequently mutated driver genes are *ACRV2A, APC*, and *TGFBR2*, which contrasts with non-hypermutant cancers where the classic triplet of *APC, TP53*, and *KRAS* are the most commonly mutated driver genes (Muzny et al. 2012). Assuming the selective pressures in these two subtypes are similar during oncogenesis, the varied driver dominance is surprising given the different molecular pathways in which these genes act.

Such comparisons across genes are echoed within genes. Accumulating evidence suggests that different driver genes and/or mutations in the same gene can have varied oncogenic strength in CRCs of the same anatomical site (Muiños et al. 2021; Haigis 2017; Temko, Tomlinson, et al. 2018; Cannataro et al. 2017; Brunet Guasch et al. 2025). For example, codon 12 mutations of the oncogene *KRAS* result in more potent biochemical activation of the K-ras protein, increased potential for oncogenesis, and elevated in-vitro growth rates, relative to other cancer-associated *KRAS* alleles (Haigis 2017). Accordingly, across cancer types which commonly have mutated *KRAS, >*90% of *KRAS* mutations are at codon 12. It is thus surprising that hypermutant CRCs with defective polymerase epsilon proofreading (*POLE-*mutant) are enriched for what appear to be functionally suboptimal *KRAS* variants outside codon 12 (Castro-Giner et al. 2015; Haigis 2017; Favre et al. 2022) (Fig. 1A). This enrichment is arguably unexpected, as the functional consequences of the different *KRAS* alleles presumably do not vary with *POLE* mutational status or underlying mutation rate or spectrum. With the clinical consequences of atypical *KRAS* alleles under active investigation (Loree et al. 2021), there is a need to understand the mechanisms leading to their presentation.

**Figure 1.**
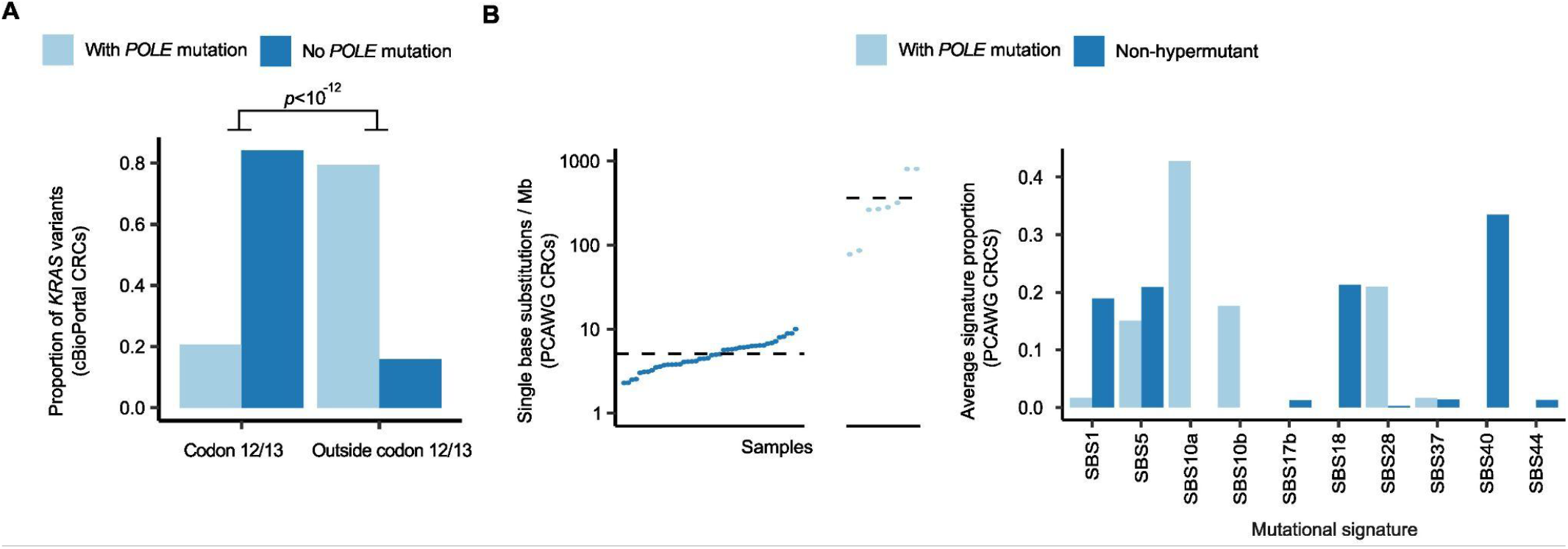
Atypical KRAS variants and mutational processes in POLE-mutant colorectal cancer. **A:** Colorectal cancers with pathogenic *POLE* exonuclease domain mutations show a significant reduction in codon 12 and 13 *KRAS* variants, which are thought to provide a strong oncogenic selective advantage (cBioPortal data, *n*=1528 CRCs without *POLE* mutation and *n=*29 *POLE*-mutant CRCs, Fisher test with odds ratio = 20.3). **B:** (Left) Single base substitution mutation burden for *POLE*-mutant colorectal cancers, and non-hypermutant colorectal cancers in PCAWG data (*n*=44 non-hypermutant CRCs and *n=*9 *POLE*-mutant CRCs). Dots are individual tumours, dashed line gives the average; (Right) Average mutational signature proportions for *POLE*-mutant and non-hypermutant colorectal cancers in PCAWG data.

Generally, a dichotomy of driver mutation classification has been suggested (Castro-Giner et al. 2015). Particular variants, such as codon 12 *KRAS* mutations, could provide an ‘near-optimal’, strong selective advantage, in comparison to weaker, ‘sub-optimal’ drivers; variant classification being determined by some combination of the observed mutation frequency and molecular or biochemical functional evidence (Haigis 2017; Temko, Tomlinson, et al. 2018; Kumar et al. 2020; Muiños et al. 2021). In particular cancer subtypes, the natural explanation for observing weaker drivers is that the active mutational processes in those cancers create a mutational bias towards these drivers. Certain cancers have defects in DNA repair or replication processes or may be repeatedly exposed to specific mutagens (Koh et al. 2021). Such effects cause excess somatic mutations to specific nucleotide changes and sequence contexts and could result in sub-optimal drivers receiving disproportionately high levels of new mutations. Moreover, the excess mutations result in an abundance of mutated driver genes (Cornish et al. 2024), which potentially shifts the epistatic landscape to favour sub-optimal drivers.

Indeed, the exposure of a tumour to mutational processes is associated with the driver variants present in the cancer at sequencing (Temko, Tomlinson, et al. 2018; Poulos et al. 2018; J. K. L. Wong et al. 2022). In keeping with this explanation, *POLE-*mutant CRCs have a distinct mutational profile, characterised by an abundance of T[C>A]T and T[C>T]G substitutions, and are associated with COSMIC mutational signatures 10a, 10b, 28 (Tate et al. 2019) (Fig. 1B). Moreover, they exhibit generally elevated mutation rates, that is a substantially increased number of mutations per cell division, resulting in orders of magnitude higher mutation burden than other cancers (Rayner et al. 2016) (Fig. 1B). The counter-argument made to the ‘mutational process’-centred explanation for weak drivers is that, as cancers develop over many years, and their clonal composition is determined by an optimising evolutionary process, cells containing sub-optimal driver variants should be displaced by fitter clones containing optimal drivers (Castro-Giner et al. 2015). On the other hand, in hypermutant cancers weak driver variants would arise more rapidly, offering a crucial head-start in the competition with nascent strong driver clones.

The questions therefore are whether mutational biases can compensate for sub-optimal selective effects, the extent of bias required, and the role played by the elevated mutation rate. In classical mathematical models of carcinogenesis (Armitage and Doll 1954) - that assume all drivers are acquired prior to tumour expansion - increases in mutation rate would be expected to accelerate tumour development, but not alter which drivers occur. However, for the arguably more realistic scenario of drivers being acquired as a neoplasm grows, such as *KRAS* in CRCs (Fearon and Vogelstein 1990), the picture is less clear.

To evaluate these arguments, in this study we employ a minimal mathematical model to investigate the conditions under which weak driver mutations acquired during tumour progression will be observed when the tumour is sampled. We consider competing clonal lineages with either strong or weak driver mutations that arise at different rates owing to variable mutational biases. We provide mathematical expressions that relate the observed driver mutation to the degree of mutational bias, the selective advantages, and the per-division mutation rates. With our mathematical results, we then examine whether the atypical *KRAS* variants in *POLE*-mutant colorectal cancers can be explained by the mutational processes operating in those tumours.

## Results

### Mathematical model of competing driver acquisition during tumour growth

Motivated by the classic adenoma to carcinoma sequence in colorectal cancer (Fearon and Vogelstein 1990), we assume tumours are initiated from a precursor cell type, for example when mutational inactivation of both *APC* alleles has occurred. While we use the terminology of proliferating cells, one may interpret ‘cells’ more broadly as the fundamental evolutionary unit in early tumourigenesis; for example colonic crypts proliferating via fission in nascent adenomatous polyps (W.-M. Wong et al. 2002).

During tumour growth, cells may acquire strong or weak driver mutations, such as codon 12/13 versus alternative driver mutations in *KRAS* (e.g. codon 117 or 146), leading to the increased proliferation of cells with those drivers. Conceptually, an increased mutation rate biassed towards weak drivers, such as that caused by *POLE*-associated mutational processes, would suggest cells with those mutations (the weak driver clone) would arise relatively quickly, resulting in dominance of the weak driver clone due to this ‘head-start’. However, over long enough time scales, by the enhanced selective advantage of the strong driver mutation, it is expected that eventually the strong driver clone will arise and sweep through the tumour. Thus, due to mutational biases, the dynamics of clone dominance would shift during growth. Which driver variants are ultimately detected when a cancer is sequenced is determined by a variety of factors including: which clones acquire further driver events, survive bottlenecks, or remain dominant at sequencing. Given reasonable biological constraints for mutational and selective parameters, our aim is to assess when mutational processes can result in the detection of non-optimal drivers through a mathematical model.

In the model (Fig. 2A), after tumours are initiated from a single founding ‘precursor’ cell, e.g. following loss of *APC*, the precursor cells divide at a baseline rate *b*_pre_ >0. When precursor cells divide, a daughter cell may stochastically acquire secondary driver mutations, either a weak driver with chance *µ*_weak_ or a strong driver with probability *µ*_strong_. As the vast majority of CRCs contain only one detectable *KRAS* variant (van Ginkel, Tomlinson, and Soriano 2023), and excessive oncogenic RAS signalling - potentially achievable through co-occurrence of both a weak and strong driver - may induce senescence (Li et al. 2018), we assume that cells only have either a weak or a strong driver, with each driver type arising only once. When comparing between cancer subtypes, the expected mutation number per division, or total driver mutation rate, *µ*_weak_+*µ*_strong_ may vary. However, even if the total driver mutation rate remains fixed between subtypes, the proportion of new mutations which are either strong or weak drivers may differ. We therefore define the ratio of the weak to strong driver mutation rate as the mutational bias to weak drivers, *β* = *µ*_weak_ / *µ*_strong_ which can, in principle, be estimated by applying the activities of measured mutational signatures to the local DNA sequence contexts of the drivers.

**Figure 2.**
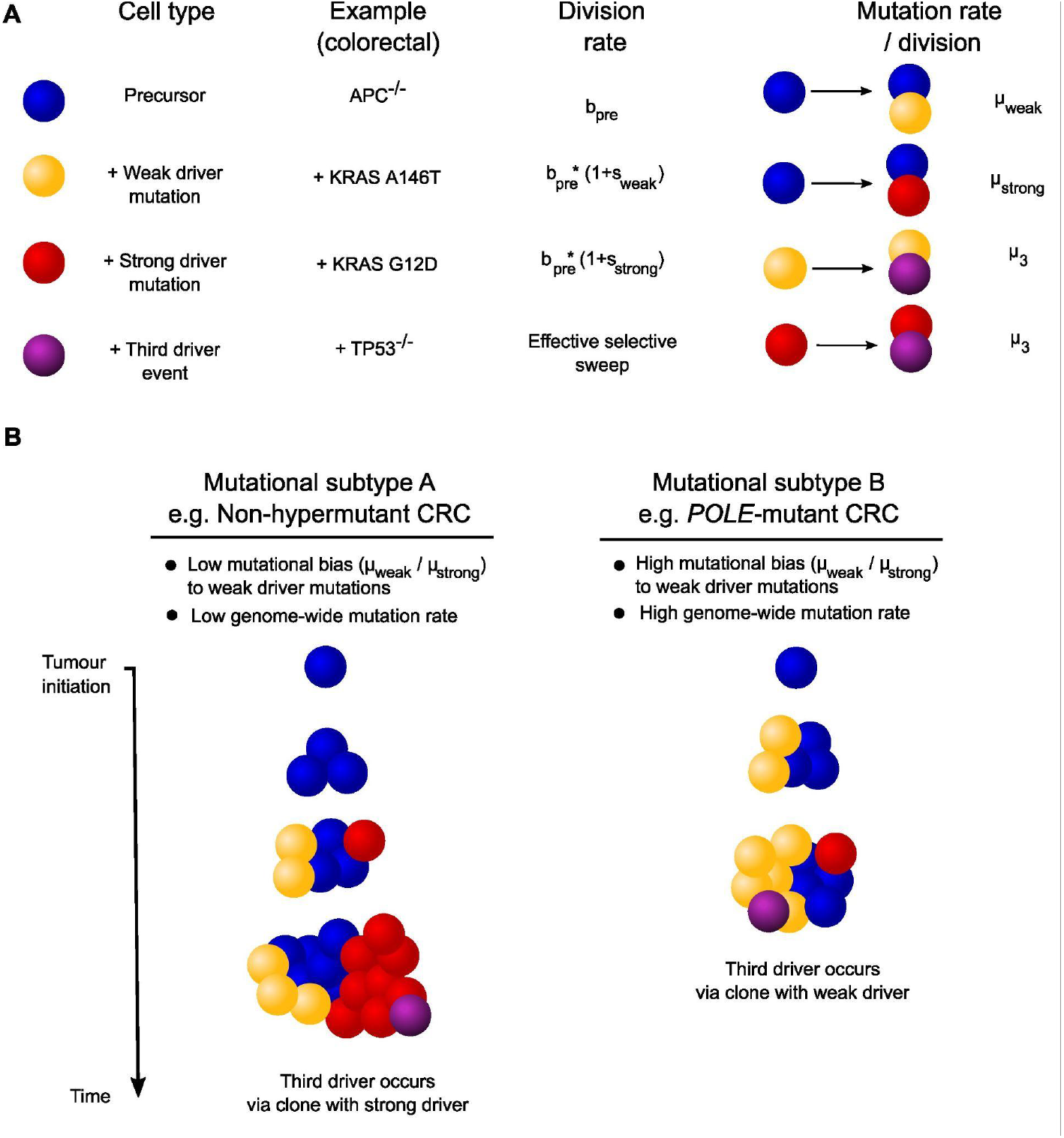
Mathematical model for variable mutational processes and selection in cancer evolution. **A:** In the model, tumours are founded with precursor cells which can randomly acquire weak or strong driver mutations at division. Weak or strong driver mutations increase proliferation rates, with strong drivers providing a larger selective advantage. Cells with secondary drivers can go on to acquire tertiary drivers, which initiates an effective selective sweep (cells without third driver are undetectable). **B**: Schematic highlighting the effect that different mutational processes can have on tumour evolution amongst cancer subtypes, without differential selection between the subtypes. In hypermutant tumours (right) due to a high mutational bias, weak driver variants arise quickly, providing a head-start to the weak driver clone which can be rapidly capitalised upon by further driver events - ultimately leading to weak drivers being detected in sequenced tumours. In contrast, for non-hypermutant cancers with a tempered mutational bias to weak drivers, the strong selective advantage provided by optimal drivers enables the strong driver clone to become dominant within the tumours.

Oncogenic *KRAS* mutations accelerate cell division rates and enhance crypt fission in the intestine (Snippert et al. 2014; Paterson et al. 2020). Motivated by these observations, we assume cells with weak or strong driver mutations divide at a rate increased by a factor of either 1+*s*_weak_ or 1+*s*_strong_ relative to precursor cells, respectively, with strong drivers providing a larger proliferative advantage, that is *s*_weak_ < *s*_strong_. Cells with either strong or weak drivers acquire a tertiary driver event at rate *µ*_3_ per division, which is assumed to initiate a rapid selective sweep, leaving other cell types undetected in any resultant cancer (although perhaps present at low frequencies). We assume that the tertiary driver is only selected if the cell has either a strong or weak secondary driver. Motivated by findings that *POLE* mutations occur prior to bi-allelic mutation of *APC (Temko, Van Gool, et al. 2018)*, we suppose that all mutation rates throughout growth can be subtype dependent.

The argument that weak drivers could be observed due to a head-start relies on there being a phase during growth such that the weak driver clone is abundant, when further oncogenic events then occur. Thus, we initially focus on which clone is dominant, i.e. comprises more than 50% of the tumour, as a function of tumour size.

### Competing clonal dynamics through tumour growth

We first investigated clonal dynamics of the model through stochastic simulations (supplementary information), starting from a single precursor cell, and tracking the emergence and growth of clones with weak or strong driver mutations. How parameter combinations determine which mutations are expected to be detectable will be mathematically analysed momentarily; here we illustrate the broad behaviour of the model and show how modest alterations to individual parameters, within biologically plausible parameter ranges (supplementary information), can considerably alter the dynamics.

Disregarding the case where mutation rates to both sets of drivers are uniformly low - which results in sustained dominance of the precursor cell type - the evolutionary trajectories of simulated tumours can be classified into three scenarios (Fig. 3A-C). Fig. 3A displays a scenario such that, due to a small weak driver mutational bias, both clones arose at similar times and thus the selective advantage of the strong driver ensured its dominance for the majority of tumour growth (i.e. before its detection or the next selective sweep). Conversely, Fig. 3B shows that simply increasing the mutational bias 50 fold resulted in a sufficient head-start for the weak driver clone to remain dominant. Thus, altering the mutational bias alone while keeping selection parameters the same is enough to alter the expected dominant driver mutation.

**Figure 3.**
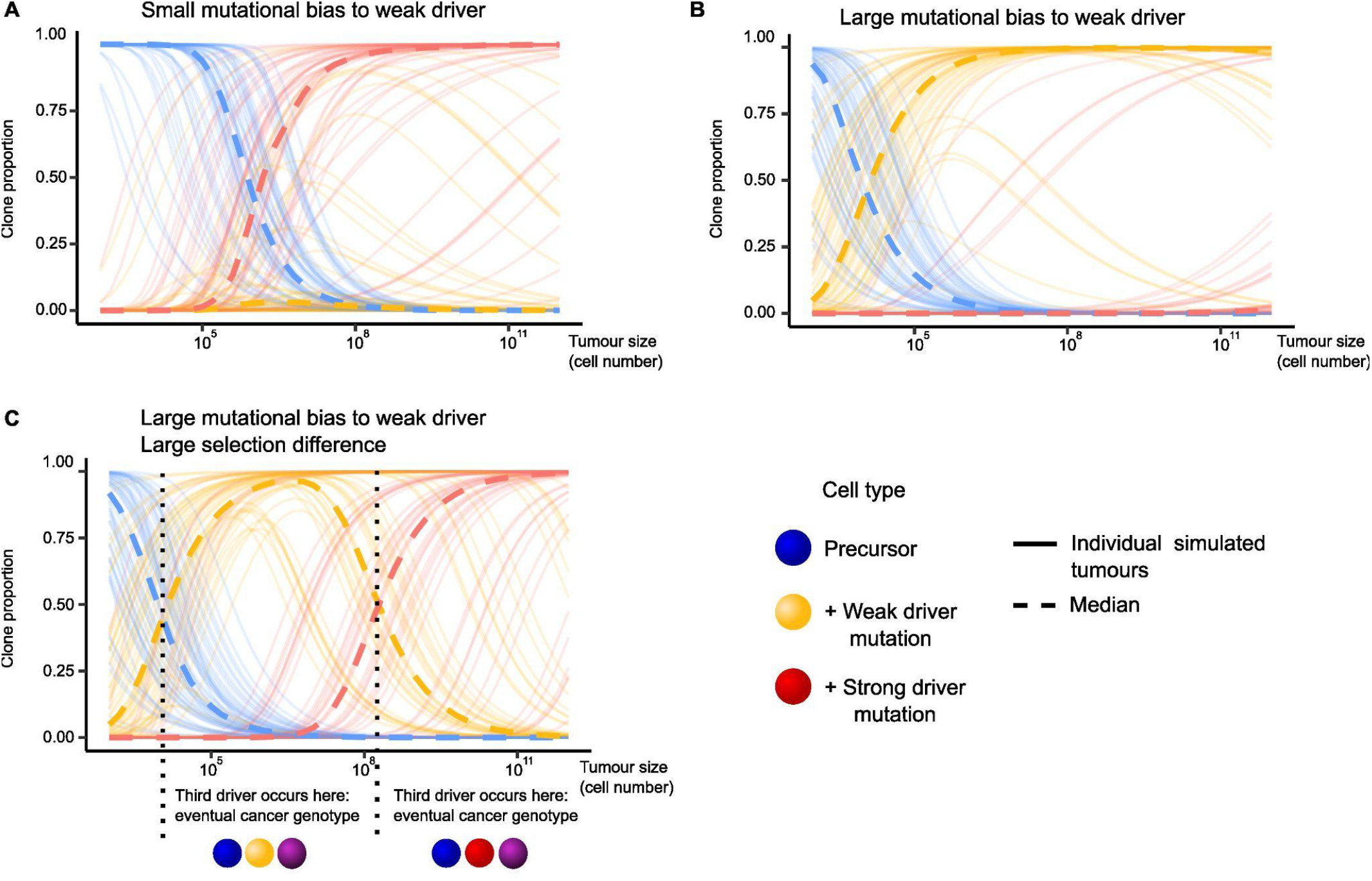
Mutational biases can lead to dominance of the weak driver clone. Stochastic simulations of the model. **A:** Similar mutation rates between weak and strong drivers results in an insufficient head-start for sustained dominance of the weak driver clone. **B:** Mutational bias increased 50 fold relative to the parameters in A, resulted in sustained dominance of weak clone. **C:** Mutation rates as in A, but with the strong driver selective advantage doubled, which resulted in a transient window of weak clone dominance; third drivers occurring within this window would lead to detection of weak secondary drivers. 50 simulations are shown per parameter set. Parameters: *b*_pre_ =1, A. *μ*_weak_ = 10^−4^, *μ*_strong_=5*10^−5^ (hence bias = 2), *s*_weak_=1.5, *s*_strong_ = 2.5; B. *μ*_weak_ = 5*10^−3^, *μ*_strong_=5*10^−5^ (hence bias = 100), *s*_weak_=1.5, *s*_strong_ = 2.5; C. *μ*_weak_ = 5*10^−3^, *μ*_strong_=5*10^−5^ (hence bias *β*= 100), *s*_weak_=1.5, *s*_strong_ = 5.

The selective sweep setting of a strong driver outcompeting the weak driver clone is shown in Fig. 3C. These simulations used the same large mutational bias as Fig. 3B but amplified the selective difference between the weak and strong drivers by simply doubling s_strong_. This led to the weak driver clone arising early due to the mutational bias, resulting in a transient window of weak driver dominance; however eventually a strong driver mutation and its lineage became fixed within the tumour. Thus, if a tertiary driver event, e.g. *TP53* mutations occurred at intermediate tumour sizes, the weak driver would be dominant subsequently in the tumour (Fig. 3C). This scenario echoes the argument that as cancer is evolutionarily optimising (Fortunato et al. 2017), the strongest variants are expected to be dominant even in hypermutated cancers.

So far we have neglected the acquisition of tertiary drivers, although it is already clear that the timing of further drivers plays a critical role. Late arriving tertiary drivers will occur in cells with strong driver mutations. In hypermutated cancers, simply due to their accelerated mutation rate, tertiary drivers could occur within the window of weak driver dominance, leading to the selection, and ultimately detection, of weak drivers. Hence, an altered mutation rate - even with the same mutational biases across the genome - offers another route such that different drivers are ultimately detected between cancer subtypes without invoking selection. To quantify the degree of bias and/or increased mutation rate required such that mutational processes can control detected drivers, we turn to a mathematical analysis.

### Mutation-selection conditions that favour a molecular pathway via a weak second driver to a tertiary driver

For a given evolutionary pathway to the third driver, the average time until the third driver occurs can be obtained analytically (supplementary information). We introduce the scaled selection difference Δ = (*s*_strong_ - *s*_weak_)/[(1+*s*_strong_)(1+*s*_weak_)] which is the difference in selection coefficients divided by the product of the fold increases in growth rate. Then, recalling that the mutational bias is *β* = *µ*_weak_ / *µ*_strong_, using branching process theory (Durrett and Moseley 2010; Durrett 2015; Nicholson et al. 2023), we find that the third driver mutation occurs earlier in the weak driver clone if

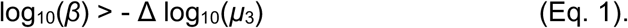

A comparison with simulation, showing that Eq. 1 correctly delineates the parameter space, is displayed in Fig. 4A. As expected, a large mutational bias to the weak driver and/or a small selective difference between drivers favours the third driver occurring within the weak driver clone, during the brief (Fig. 3C), or extended (Fig. 3B), period of weak driver clone dominance.

**Figure 4.**
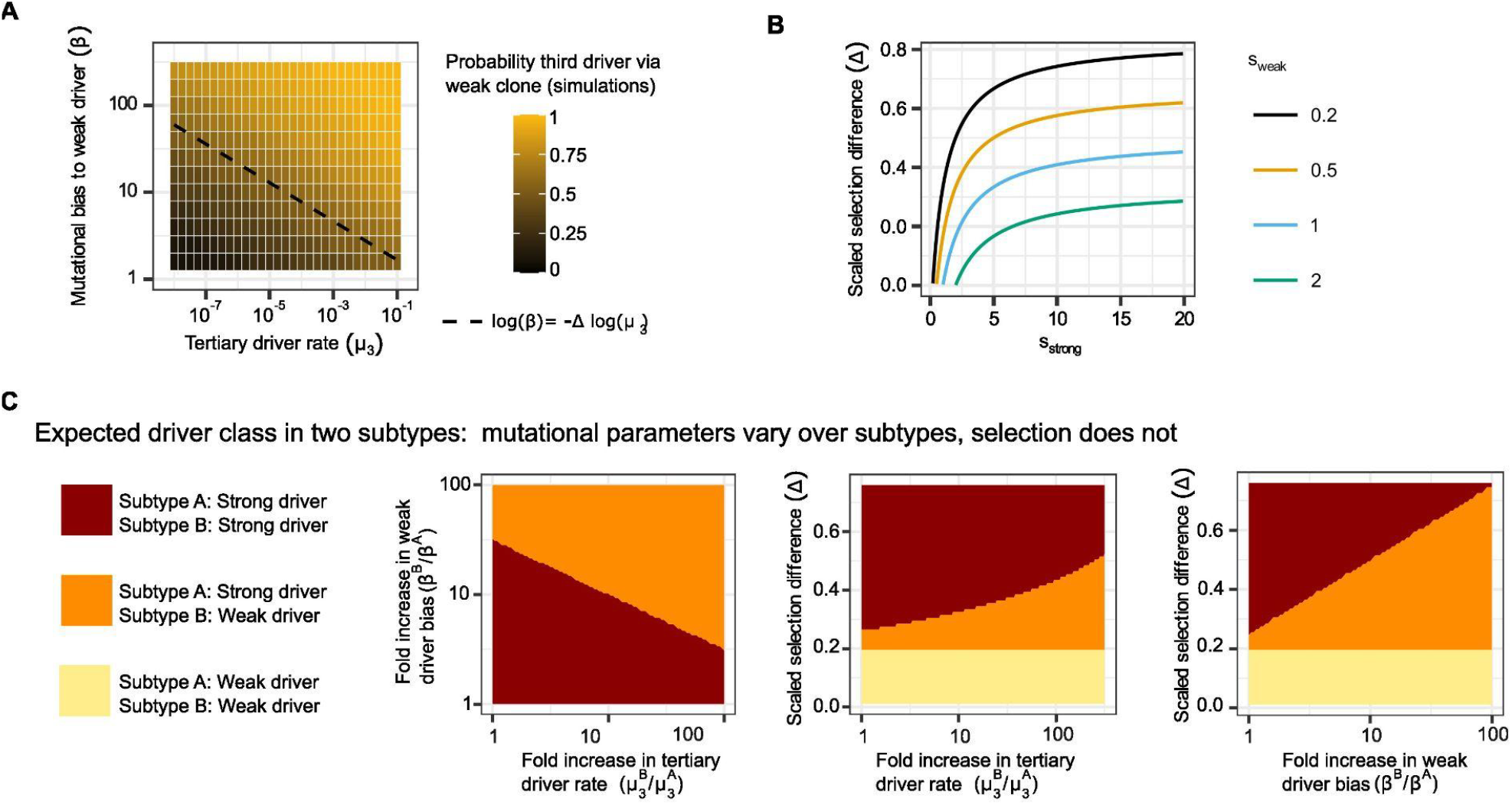
High mutational biases and tertiary driver rate result in the third driver occurring via weak driver clone. **A:** Varying the weak driver mutational bias and the mutation rate to acquire a third driver, the proportion of 100 simulations that resulted in the third driver occurring via the weak driver clone is displayed. Dashed line given by Eq. 1. Fixed parameters: *b*_*p*_ =0.2, *µ*_weak_ *=* 5*10^−3^, *s*_weak_=0.5, *s*_strong_*=*1.25. **B:** The scaled selection difference, Δ = (*s*_strong_ - *s*_weak_)/[(1+*s*_strong_)(1+*s*_weak_)], as a function of the selection parameters; Δ controls the mutational bias and tertiary driver rate required such that it is more likely the tertiary driver is acquired within the weak driver clone - specifically, this occurs when log_10_(*β*)/log_10_(*µ*_3_^-1^) is above the displayed line. **C:** Comparing two hypothetical cancer subtypes A and B with different mutational biases and overall mutation rates, we show parameter regimes where the evolutionary path to a third driver is expected to be the same or different between the subtypes, that is via a weak or strong second driver. Regimes determined via Eq. 1. Fixed parameters: (left) *β*^A^= 10, Δ=0.5, *µ*_3_^A^=10^−5^; (centre) *β*^A^= 10, *β*^B^=20, *s*_weak_=0.3, *µ*_3_^A^=10^−5^; (right) *β*^A^= 10, *s*_weak_=0.3, *µ*_3_^A^=10^−5^, *µ*_3_^B^=10^−4^.

Notably, modulation of the tertiary driver rate, *µ*_3_, while keeping all else equal, is sufficient to alter whether tertiary drivers occur within the weak or strong driver clone. If further driver events occur relatively quickly after a secondary driver clone appears, as might be the case in hypermutant cancers, this would favour the weak driver evolutionary path, ultimately resulting in detectable weak drivers in those cancers. In contrast, only the mutational bias to the weak driver determines the evolutionary path taken to the third driver, not the absolute values of *µ*_weak_ and *µ*_strong_. Explicitly, if we consider a specific gene with 16 weak driver sites and 2 strong driver sites, and a uniform per-base mutation rate, then the left-hand-side of Eq. 1 is always log_10_(8) regardless of the numerical value of the per-base mutation rate. The scaled selection difference, Δ, controls the degree of bias required and takes values between 0 and 1 (Fig. 4B). When the selection coefficients are small, Δ ≈ *s*_strong_ - *s*_weak_, and so in this setting, linear increases to *s*_strong_ require exponential increases to *β* or *µ*_3_ to ensure the weak driver pathway is fastest, demonstrating the dominant force of selection. At the other extreme, with a highly potent strong driver so that *s*_strong_ is large, Δ ≈ 1/(1+ *s*_weak_) (right-limit in Fig. 4B), the weak driver path may still be more likely, as long as the third driver occurs rapidly to capitalise on the weak driver clone’s limited head-start.

Motivated by different drivers being present in CRC mutational subtypes, we next investigated when modulation of mutational processes alters whether a tertiary driver occurs via the weak or strong clone. Suppose we are considering CRC subtypes A and B, and for a fixed set of weak, strong, and tertiary drivers, these subtypes have associated mutational biases and tertiary driver rates, *β*^A^ and *µ*_3_^A^, and, *β*^B^ and *µ*_3_^B^, respectively. We assume that the selective effects of each driver mutation, whether weak or strong, do not vary with subtype.

We mapped changes in mutational processes between the subtypes to whether the same driver is to be expected using Eq. 1 (Fig. 4C) and found that increasing the relative tertiary driver mutation rate between the subtypes, *µ*_3_^B^/*µ*_3_^A^, can be sufficient for different expected evolutionary paths occurring. A high relative tertiary driver rate could occur simply by subtype B having a genome-wide higher mutation burden per cell division, as might be expected in the hypermutant setting. Altering the relative tertiary driver rate was able to alter the evolutionary path, even when the mutational bias was equal between the subtypes, i.e. *β*^A^ = *β*^B^ (see bottom right corner of left panel in Fig. 4C). This effect would not be present in classical mathematical models of tumourigenesis that neglect clonal growth (Armitage and Doll 1954), where simply increasing the total mutation burden would accelerate mutation accumulation but would not alter the likelihood of one mutation being observed over another.

Interpreting the *KRAS* variants observed in hypermutant cancers through this minimal model, these results suggest that a combination of the altered mutational bias (a higher propensity for mutations occurring at given genomic contexts) and the increased genomic per-division mutation rate are sufficient to explain the observed non-optimal *KRAS* variants. To explicitly address this, we next classify drivers as weak or strong to obtain numerical values for the mutational biases.

### *KRAS* drivers in *POLE*-mutant cancers

A high frequency of unusual *KRAS* mutations has previously been noted in *POLE-*mutant tumours (Castro-Giner et al. 2015). To quantify this effect, we analysed publicly available CRC mutation data (1557 independent samples obtained from cBioPortal, see Methods). *POLE-*mutant CRCs - classified by the presence of a non-conservative missense mutation within the exonuclease domain of *POLE* - showed a significantly altered distribution of mutated *KRAS* codons compared to CRCs without a *POLE* mutation (Fig. 5A). Exemplifying the difference, 17% of *POLE-*mutant cancers carried a codon 117 *KRAS* variant, compared to only 1% of cancers without a *POLE* exonuclease domain mutation.

**Figure 5.**
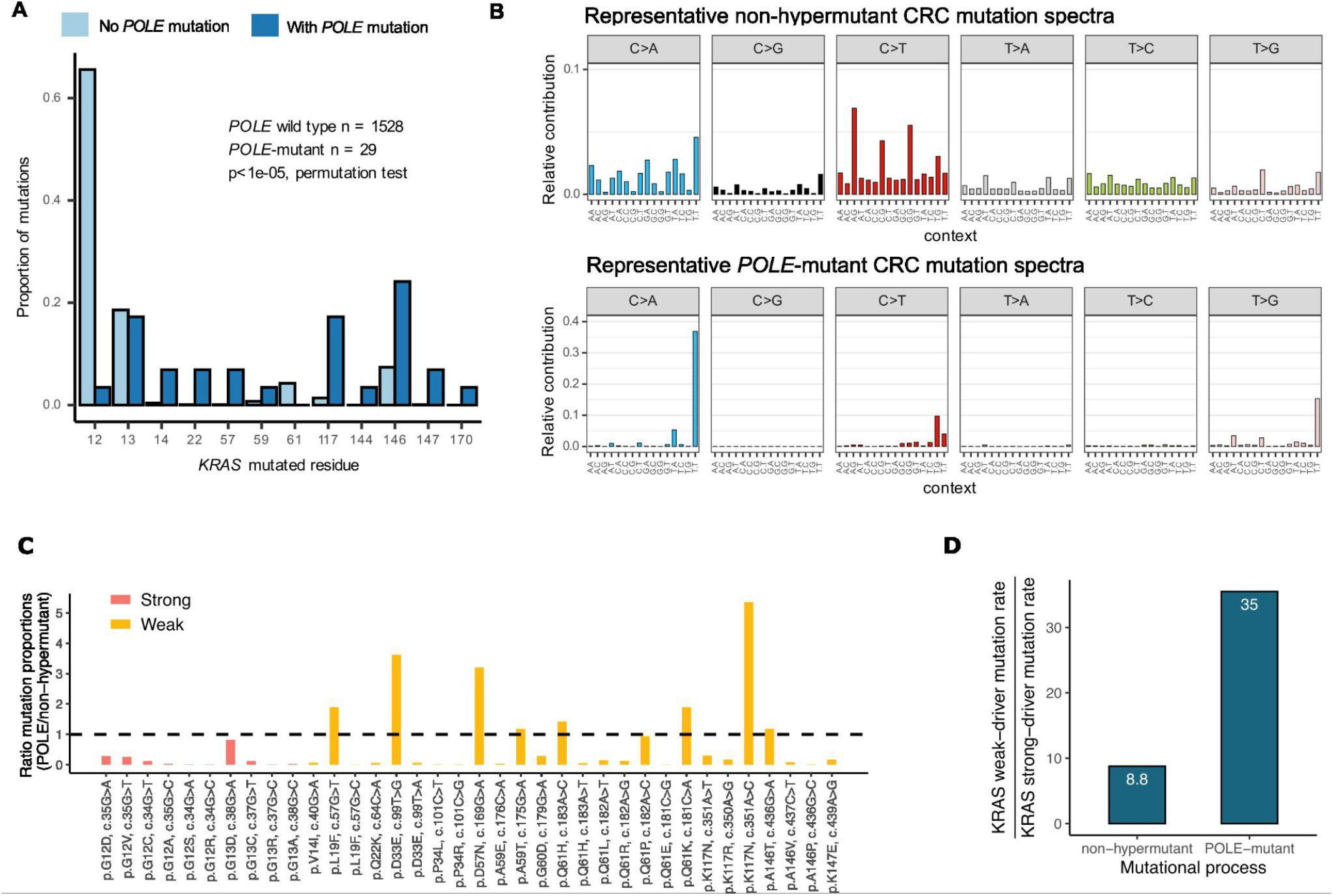
Altered mutational processes and atypical KRAS variants in POLE-mutant CRCs. **A**: *POLE*-mutant CRCs display a significantly different distribution of *KRAS* mutations compared with wild-type *POLE* tumours. **B**: Representative mutation spectra for single base substitutions for *POLE*-mutant and non-hypermutant CRCs. **C:** Ratio of the proportion of mutations expected to fall at specific *KRAS* variants under *POLE*-mutant mutational spectra or non-hypermutant CRC spectra. Here, variants in codon 12 or 13 were classified as strong drivers, other variants as weak. **D:** Mutational bias to weak versus strong *KRAS* variants (classified as in **C**) under *POLE* associated mutation spectrum or non-hypermutant CRC mutational spectrum

A curated list of *KRAS* colorectal cancer driver variants was considered based on multiple data sources including intOGen (Martínez-Jiménez et al. 2020), cBioPortal (Cerami et al. 2012), COSMIC and UK 100,000 Genomes Project (Cornish et al. 2024) DNA sequencing, in addition to annotation from functional studies. Varied criteria were used to classify *KRAS* driver variants (supplementary table 2) into weak or strong drivers including: the bioactivation score defined by in-vitro measures of allele-specific protein biochemistry (Haigis 2017); assigning variants in codons 12 or 13 as strong drivers which was inspired by their prior classification as canonical *KRAS* variants (Loree et al. 2021); and variants observed at disproportionately high levels in non-*POLE* mutated CRCs when assuming variant frequency is controlled only by mutational signatures. For each of the criteria used to classify the drivers, there was a significant (Fisher’s test, *p<*0.05, Supplementary Fig. 1) enrichment of weak drivers in *POLE--*mutant CRCs compared to CRCs without a *POLE* mutation. For example, taking variants in codon 12 or 13 to be strong drivers, and other *KRAS* variants as weak, 74% of *KRAS* driver variants in *POLE--*mutant CRCs were weak drivers compared to only 15% in CRCs without a *POLE* mutation. This enrichment was preserved upon stratifying tumours by anatomical location (distal versus proximal, Supplementary Fig. 1).

*POLE*-mutant cancers have distinct mutational signatures (Rayner et al. 2016). To assess how the signatures translate into local mutational biases to different *KRAS* sites, representative single-base-substitution mutation spectra were constructed for both *POLE--*mutant CRCs and standard (non-hypermutant) CRCs using signature data reported by the PCAWG project (Campbell et al. 2020). For the standard CRC spectrum, we averaged the mutation spectra of the non-hypermutant CRC samples, while to define the representative *POLE-*mutant spectrum, we mixed COSMIC signatures 10a, 10b, and 28 according to the average proportions found in the *POLE-*mutant CRC samples (Fig. 5B).

We estimated the proportion of new mutations expected to result in each *KRAS* variant under both representative spectra. Analysing the ratio of mutation proportions (Fig. 5C), highlighted c.351A>C p.K117N, which is consistently labelled as a weak driver and has a ∼5.4 fold increase in mutation proportion under the *POLE*-mutant spectrum compared to the standard CRC mutation spectrum, and c.35G>A p.G12D, which was consistently classified as a strong driver and has ∼3.5 fold decrease in mutation proportion under the *POLE*-mutant spectrum. For 28/36 of the *KRAS* variants, the estimated mutation proportion was higher under the standard CRC spectrum, as a consequence of the flatter spectrum.

Across the driver classification criteria, the estimated mutational bias to the weak drivers compared to the strong drivers, *β* = *µ*_weak_/*µ*_strong_, was greater than 1 under both representative spectra. Focusing on the codon 12/13 strong driver criterion, *β* = 35 under the *POLE*-mutant spectrum whereas *β* = 8.8 using the standard CRC spectrum. Thus, under mutational spectra considerations alone, 35 times as many weak drivers compared to strong drivers are expected in *POLE*-mutant cancers, while 8.8 times more weak drivers are expected in non-hypermutant CRCs. In part, this was simply due to classifying more drivers as weak compared to strong, of the 36 *KRAS* driver variants considered, only 8 are in codon 12 or 13. However, a uniform mutation rate over variants would result in *β* = 28/8 = 3.5, illustrating the effect of the mutational processes. Notably, under each criterion the mutational bias to weak drivers was larger under the *POLE* mutational spectrum than the standard CRC mutational spectrum, ranging from a 2.25-fold to 5-fold increase. In particular, an approximately 4-fold (35/8.8) increase was estimated under the codon 12/13 criterion.

We used the mathematical model to survey conditions such that the increased bias to weak *KRAS* drivers as estimated in *POLE*-mutant cancers, in combination with an increased per-division mutation rate, would lead to observing *KRAS* weak drivers (Fig. 6). Due to the vastly increased mutation burden in *POLE-*mutant cancers, we assumed that all mutations generated during tumour growth in *POLE-*mutant cancers followed the *POLE-*mutant spectrum, while mutations generated in non-*POLE* cancers were assumed to follow the standard CRC spectrum. For illustration, we took the mutational biases obtained from classifying *KRAS* variants based on the codon 12/13 criterion, Fig. 5D. Using the other classification criteria gave similar results (Supplementary Fig. 2). Further, we assumed that *POLE* cancers have a 100X fold increase in mutation rates.

**Figure 6.**
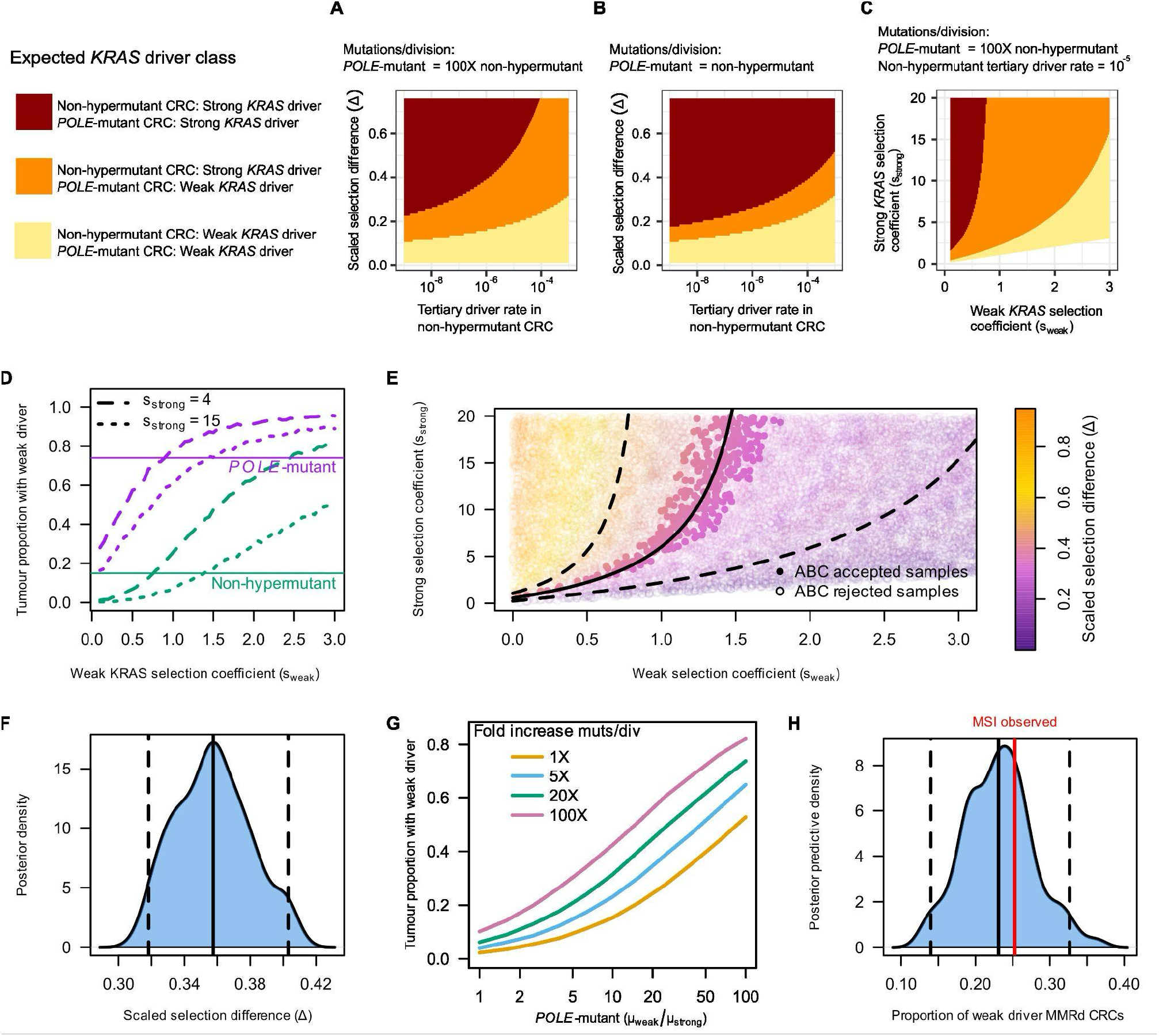
Differences in mutational processes can alter KRAS drivers between non-hypermutant and POLE-mutant colorectal cancers. **A:** Heatmap displaying parameters such that the expected pathway to the third driver differs or remains the same between non-hypermutant and *POLE*-mutant CRCs, using estimated mutational biases to weak and strong *KRAS* drivers (Fig. 5D). **B:** As in **A**, with equal mutation rates between subtypes. **C**: As in **A**, assuming the tertiary driver rate in non-hypermutant CRCs = 10^−5^. **D:** Simulated tumour proportion with weak *KRAS* drivers. Horizontal lines display the proportion observed in the cBioPortal cohort. **E:** Prior parameter samples used to simulate tumour cohorts. Parameters resulting in simulations similar to observations, i.e. posterior samples, are opaque. Dashed lines indicate bounds on selection coefficients suggested by Eq. 1 (as in **C**), filled line indicates relationship between coefficients with Δ=0.36, the posterior mean. **F:** Posterior distribution of scaled selection difference. **G:** With ABC optimal selection parameters, simulated tumour proportion with weak drivers vs weak mutational bias. Fold increase in mutation rate is relative to non-hypermutant value, non-hypermutant weak bias = 8.8. **H:** With ABC posterior parameters, estimated MSI CRC mutational bias to weak drivers, and 10X fold increase in mutation rate per division, the density of tumour proportion with weak drivers in simulated cohorts is shown. The vertical red line displays the observed proportion. Estimated mutational biases to drivers as in (Fig. 5D) used throughout. Other parameters: (A, B) *s*_weak_=0.3, (C, E, F, G, H) non-hypermutant *µ*_*3*_ *=*10^−5^, (D) non-hypermutant *µ*_weak_=10^−7^, non-hypermutant *µ*_*3*_ *=*10^−5^.

The expected pathway to the third driver in both *POLE* and non-hypermutant CRC subtypes is displayed in Fig. 6A, obtained by evaluating Eq. 1 while ranging over the tertiary driver mutation rate in non-hypermutant CRCs, *µ*_3_, and the scaled selection difference, Δ. A considerable subset of the plausible parameter space resulted in *KRAS* weak drivers being observed in *POLE*-mutant CRCs, with *KRAS* strong drivers being observed in non-hypermutant CRCs. This space is markedly reduced if the *POLE* mutation does not increase the tertiary driver mutation rate (Fig. 6B). To further examine the role of selection, we fixed a representative value for the tertiary driver rate in non-hypermutant CRCs as *µ*_3_=10^−5^ (Bozic et al. 2010). In Fig. 6C, a wide range of selection coefficients is shown to be compatible with the different *KRAS* variants observed in the different CRC subtypes. Therefore, at a broad level, within the mathematical model, the different *KRAS* driver variants observed in *POLE*-mutant CRCs relative to non-hypermutant CRCs can be explained by the altered mutational processes between these cancer groups without invoking differential selection pressures.

We queried whether comparing model output with the observed tumour proportions carrying weak drivers held further information on the parameters which best explained the data. Focusing on the codon 12/13 strong driver criterion, via initial simulations we found that parameters that resulted in matching the empirically observed proportions depended jointly on *s*_weak_ and *s*_strong_ (Fig. 6D). For parameter inference, we performed an approximate Bayesian computation (ABC) analysis, with parameter priors on *µ*_weak_ for non-hypermutant CRCs and *s*_weak_ and *s*_strong_; all other parameters were fixed as detailed above. Posterior samples confirmed the dependency of *s*_weak_ and *s*_strong_ (Fig. 6E). Mimicking the lack of dependency on absolute rates in Eq. 1, the posterior for non-hypermutant *µ*_weak_ was uninformative (Supplementary Fig. 3). In contrast, the posterior distribution of the scaled selection difference Δ was unimodal (mean = 0.36, 95%CI = (0.32, 0.40), Fig 6F) and well explained the dependency in the selection coefficients (filled line, Fig 6E). Assuming *KRAS* variants increase crypt fission, the posterior selection coefficients imply that the codon 12/13 variants provide an average of 5.56 (CI95=2.37 to 8.46) fold greater increase in fission rate relative to the weak drivers, i.e. (1+*s*_strong_)/(1+*s*_weak_). The posterior selection samples comprised a considerably smaller area of the parameter space compared to that implied by Eq. 1 (Fig. 6E), confirming the benefit of fitting against the tumour proportions.

For the optimal parameters identified in the ABC analysis, the relative effects of mutational bias and variations in mutation rate are exemplified in Fig. 6G. Under classical models of tumour initiation, while varying the mutational bias would alter the tumour proportion with weak drivers, the amplifying effects of increased mutation rate would be absent. Here, this effect is pronounced: assuming a fixed *POLE* mutational bias of *β* = *µ*_weak_/*µ*_strong_=25, the proportion of hypermutant CRCs with weak drivers increases from 28% to 61% as the fold increase in mutation rate ranges between 1 to 100 (Fig. 6G).

### Mismatch repair deficient tumours

Mismatch repair deficient (MMRd) CRCs represent an intermediate mutation rate between non-hypermutant colorectal cancers and those with *POLE* proofreading deficiency, with an approximately 10X increase in single base substitutions relative to non-hypermutant tumours (Timmermann et al. 2010; Cornish et al. 2024). Thus far we have not focused on MMRd CRCs as, while *POLE*-mutant CRCs are likely to follow the classical adenoma to carcinoma pathway, a substantial proportion of MMRd CRCs are thought to develop with activating MAPK mutations, most commonly in *BRAF*, and via the serrated polyp pathway (Nguyen et al. 2020). In these non-classical pathways, the explanatory power of our model would be diminished.

A subset of MMRd CRCs are likely to arise from adenomas and these are associated with mutant *KRAS (Jass 2004)*. We queried whether the weak driver model would be followed within MMRd *KRAS*-mutant CRCs. Given the more modest increase in mutation rate, we would predict only a modest enrichment in weak drivers relative to non-hypermutant tumours. Thus to boost our power, in addition to the cBioPortal data analysed thus far, we included 816 CRCs with oncogenic *KRAS* mutations from the 100,000 Genomes Project (Cornish et al. 2024; “National Genomic Research Library, Genomics England (2024),” n.d.), previously annotated as microsatellite stable or microsatellite unstable (MSI). Adopting the codon 12/13 strong driver criterion, tumours without a pathogenic *POLE* mutation, but annotated as MSI (likely due to MMRd), exhibited an intermediate number of weak drivers: 25% contained drivers outside of codon 12/13, a significant increase compared to the 15% observed in non-hypermutant tumours (p<10^−3^, OR = 0.51, Fisher’s test).

To assess whether these data were consistent with our model, we used previous estimates for a representative CRC MSI mutational signature (Brunet Guasch et al. 2025). This resulted in a mutational bias to weak drivers of only 6.73, a reduction compared to the 8.8 estimated for non-hypermutant CRCs. Naively this would lead to an expected depletion of weak drivers in MSI tumours. However, simulating tumour cohorts of equal size to the data, using the posterior samples from the ABC inference above, and with a 10 fold increased mutation rate in MSI tumours, led to simulated outcomes that were consistent with the observed weak driver proportion (Fig. 6H). Thus, *KRAS* acquisition in these MSI tumours is consistent with our model and parameters inferred from non-hypermutant and *POLE*-mutant tumours.

## Discussion

### Summary of findings

Our mathematical analyses provide an explanation for the apparent paradox that some cancers, especially colorectal cancers (CRCs) with hypermutator phenotypes and accompanying irregular mutational biases, tend to acquire atypical, presumed selectively sub-optimal, driver mutations in genes such as *KRAS*. Our arguments centre on a mathematical model of competing clonal lineages in nascent tumours, with precursor cells, e.g. bi-allelic *APC* mutant colorectal cells, acquiring weak or strong secondary driver mutations en route to advanced cancer. In some scenarios, we find that the strong driver clone predominates from an early stage of tumourigenesis, whereas in others, in the absence of a subsequent driver mutation, the weak (atypical) driver dominates temporarily, until the strong driver arises and displaces the weak driver in an effective selective sweep. These scenarios represent the “classical” view that natural selection in cancer evolution is optimising (Fortunato et al. 2017). However, for a wide range of biologically plausible parameter settings we predict that the weak driver would be predominant in sequenced hypermutant tumours, as observed in reality. Echoing the ‘leapfrog effect’ of evolutionary genetics (Gerrish and Lenski 1998), the weak driver predominance is facilitated via a head-start which can be capitalised upon by rapidly occurring subsequent driver events.

In line with expectations, the presence of an atypical driver is favoured by (i) a strong mutational bias towards the nucleotide context in which the atypical driver occurs and (ii) only small differences in the selective advantages of the typical and atypical drivers. A third, less expected, factor that favours weak driver detection is that the time to occurrence of the next driver mutation (*e*.*g. TP53* in CRCs) is reduced in hypermutant cancers. This effect, which is a consequence of elevated mutation rates, is not present in standard models of tumourigenesis that neglect clonal expansions prior to cancer initiation (Armitage and Doll 1954; Tomasetti et al. 2015; Temko, Tomlinson, et al. 2018), where mutation acquisition would be accelerated but the distribution of resultant genotypes unchanged. Driver mutations in CRCs with defective DNA polymerase proofreading (*POLE-*mutant) are consistent with the mutational profile associated with *POLE* deficiency (Temko, Van Gool, et al. 2018), supporting the accelerated accumulation of further drivers in these tumours.

Focussing our analysis on *KRAS* variants in *POLE*-mutant CRCs, we estimated an elevated mutational bias to weak *KRAS* drivers in *POLE*-mutant CRCs compared with non-hypermutant tumours. This elevated bias held across varied classifications of weak and strong drivers. Integrating these estimates with our mathematical model, we found that the bias, in combination with the ∼100 fold increased mutation rate found in *POLE*-mutant CRCs, is sufficient to explain the atypical *KRAS* variants observed in these hypermutant tumours. Thus, differential selective effects of driver mutations in hypermutant and non-hypermutant cancers need not be invoked to explain the atypical drivers observed in *POLE*-mutant colorectal cancers.

### Alternative hypotheses for atypical *KRAS* variants

A natural alternative hypothesis is that driver mutations have subtype-specific selective effects, i.e. weak drivers conferring a fitness advantage specifically within hypermutant cancers. While we cannot rule out this scenario, it is unclear how it would mechanistically occur in the initial stages of tumourigenesis, as: (i) we expect that the *KRAS* variants we focus on will have the same biochemical impact regardless of hypermutant status; (ii) these mutations are acquired early in colorectal tumourigenesis, and thus as similar tumour microenvironments are expected, we would predict similar phenotypic effects.

During tumour progression the selective landscape may change due to the increased mutation rate in hypermutant cancers. Indeed, hypermutants are likely more immunogenic (van Gool et al. 2015), which could reduce the advantage of driver mutations once the tumour is immunosurveilled. Recent evolutionary analyses (Tilk et al. 2022) suggest that hypermutant cancers have reduced efficiency of selection; under this model, later drivers will occur in cells with increased deleterious mutation load. In both scenarios - immunogenicity and deleterious mutation accumulation - the net effect is that the advantage of early arriving weak drivers would be even further elevated in hypermutant cancers, exacerbating the role of a weak driver head-start. If either of these general selection mechanisms occur, we would predict that the reduced time to acquire the third driver mutation in hypermutant cancers would play an even greater role.

Caveating this prediction, a specific explanatory mechanism acting independently of accelerated mutation is variable antigenicity of the different driver variants (Bear et al. 2021). In pancreatic adenocarcinomas, a trade-off between functional fitness and immunogenicity has been reported for *KRAS* alleles (Hoyos et al. 2022). Speculatively transferring this trade-off to the colorectal setting, one would predict that strong *KRAS* alleles are more immunogenic and thus may be less likely to progress to advanced cancer on a *POLE*-mutant background due to enhanced immunosurveillance. Future immunological studies comparing hypermutant adenoma and advanced carcinoma should inform on this mechanism.

Generally, epistasis is a possible cause of the pervasiveness of weaker driver genes (Castro-Giner et al. 2015). Numerous variants are likely acquired between the *POLE* and *KRAS* mutations, which could alter the sweet spot for oncogenic RAS signalling (Li et al. 2018) such that canonical *KRAS* drivers are no longer maximally oncogenic. Indeed, inferred epistatic interactions vary by *KRAS* allele (Cook et al. 2021), thus *POLE* mutational processes could bias mutations towards genes that positively epistatically interact with atypical *KRAS* alleles and antagonistically with canonical alleles. However, this hypothesis is currently challenging to evaluate as the posited interactions are at the earliest stages of tumourigenesis, whereas the cancer genomic data informs on mutation patterns at sampling. One manifestation of this mechanism is multiple weak RAS pathway drivers cumulatively acting to initiate tumourigenesis. Casting doubt on the universality of the epistatic mechanism, recent analyses (Woolley et al. 2025) found that the vast majority of CRCs with atypical *KRAS* variants did not have further driver mutations in major RAS oncogenes, and those that did were not explained by hypermutation

### Limitations

Our analysis is principally designed to examine whether the observation of atypical drivers can be explained by known factors such as mutation rates and spectra without adopting varied selection landscapes in hypermutant tumours. It has general simplifying assumptions, and some specific limitations. We have compared two scenarios in which tumour growth starts with bi-allelic *APC* mutation and in one of which a *POLE* mutation is also assumed to have occurred. While there is evidence that the *POLE* mutation occurs first (Temko, Van Gool, et al. 2018), it could be that in a subset of cases *POLE* mutations occur late in tumourigenesis, in which case no bias would be expected for early drivers, negating the explanatory power of our model.

As is common (Luebeck and Moolgavkar 2002; Paterson et al. 2020), we stop our model after a defined number of drivers have been accumulated. This is for modelling simplicity, but could be motivated by the third driver providing a large fitness advantage or that the third driver impacts the fitness of cells without it. Importantly, as we inferred a larger mutational bias to weak *KRAS* drivers relative to strong drivers in both non-hypermutant and hypermutant CRC, it is more likely that weak variants arise first in both CRC mutational subtypes. Thus, if we stopped the model after the second driver was acquired instead of the third, weak *KRAS* drivers would dominate in both CRC subtypes, in contrast with the empirical observations. It has been recently estimated that *KRAS* mutation may be the initiating event in up to a third of CRCs (Paterson et al. 2020); in such a setting, based on inferred mutational biases, we’d again expect weak variants to predominate in both tumour types.

We have adopted the simplest, branching process model of competing clonal lineages, in part so that we can compare with evolutionary parameters obtained using a similar framework. Notably we have omitted both spatial structure and immune response in our model (Noble et al. 2022; Lakatos et al. 2020). Both effects would alter the range of biological parameters consistent with differential *KRAS* alleles appearing in the non-hypermutant and hypermutant subtypes, as well as altering the likelihood of an effective selective sweep which our model assumes occurs after the third driver event occurs (Stein et al. 2026).

A general modelling limitation is the degree to which our model reflects those rare tumour trajectories resulting in detected and sequenced cancers. Cancers that present clinically may represent the right-hand tail of the distribution of net growth rates, with mutational events occurring earlier than would be indicated by somatic mutation rate estimates. Incorporating this effect is challenging as the number and type of drivers that lead to detectable CRCs is still under active investigation (Cornish et al. 2024). However, we speculate that the net effect is a heightened relevance of large selective effects and absolute mutational parameters. As the plausibility of our explanation for differential CRC drivers principally relies on relative mutational biases over genomic loci and relative mutation rates between CRC subtypes, and holds within large regions of parameter space, we expect the primary mechanistic hypothesis proposed here to retain substantial explanatory power despite this limitation.

### Conclusion and implications

In conclusion, using mathematical modelling we have shown that the null hypothesis that irregular mutational processes, in combination with the evolutionary dynamics of tumourigenesis, are sufficient to explain the enrichment of atypical mutations in hypermutant cancers.

The relation between weak or strong drivers, and non-hypermutant and hypermutant CRCs, is not one-to-one. This is exemplified by the enrichment of *BRAF* p.Val600Glu mutations - a canonical strong driver - in mismatch repair deficient (MMRd) CRCs compared to microsatellite stable CRCs (Cornish et al. 2024). *BRAF* p.Val600Glu arises from one of the least active mutational channels in MMRd CRCs (Temko, Tomlinson, et al. 2018), while a variety of atypical driver mutations can arise from more active MMRd channels. It may be that in many cases, the p.Val600Glu clone dominates or overtakes an atypical driver clone, but there could be a sufficient window for the latter sometimes to acquire a tertiary driver first and thus persist. Given that each driver oncogene generally has more sites at which atypical drivers can occur rather than typical drivers, we hypothesise that the effect of the head-start advantage could occur in a minority of cancers in the absence of hypermutation.

The observed frequency of a cancer driver, after controlling for mutational likelihood, is a common proxy for the selective strength of drivers (Temko, Tomlinson, et al. 2018; Cannataro et al. 2017; Martincorena et al. 2017). Our finding that accelerated mutation acquisition can alter which drivers are ultimately detected, even with unchanged mutational biases and selective coefficients, suggests that caution should be used in comparing driver frequencies across cancers with highly disparate mutation rates. This finding supports separate analysis of the selective landscape of hypermutant tumours (Martincorena et al. 2017; Cornish et al. 2024).

In principle, the scenario of selective sweeps rapidly being achieved via first acquiring weak drivers could repeat multiple times, leading to “polygenic” tumourigenesis involving multiple weak drivers.

There is evidence to support this view, for example that hypermutant CRCs have an increased number of driver mutations and multiple atypical driver genes (Rayner et al. 2016). The clinical impacts of atypical *KRAS* alleles is under active investigation (Loree et al. 2021; Navarro-Jiménez et al. 2025); generally whether tumours as a whole are less fit owing to the atypical mutations is currently uncertain.

## Methods

### Details of mathematical model, analytical derivations, plausible parameter range, and approximate Bayesian computation

See supplementary information.

### Mutated KRAS residues in public data

Colorectal cancer samples available on cBioPortal were downloaded (supplementary table 1), and filtered to obtain unique patient-mutation pairs. Samples were classified as *POLE*-mutant if they contained pathogenic variants in the exonuclease domain of *POLE -* comprising those variants listed in tables 1 and 2 of (Rayner et al. 2016) plus the non-conservative variants F367C, E318K, N363D, D275G, E318K. For all samples, we collated *KRAS* missense mutations. For the MSI analysis, we used data from the 100,000 Genomes Project. We used all *KRAS* mutations annotated as oncogenic, and used mutational subtype annotations (MSS, *POLE*, MSI), as specified by (Cornish et al. 2024).

### Representative mutation spectra and KRAS variant rate

Simple somatic mutation files for colorectal cancers in the PCAWG study (Campbell et al. 2020) were obtained from the ICGC (Zhang et al. 2019). Samples with pathogenic *POLE* variants were filtered for, which resulted in nine samples. For these nine samples, we downloaded the PCAWG-reported signature exposure vectors (www.synapse.org/#!Synapse:syn11804040). The POLE-associated mutational signatures 10a, 10b, and 28, defined according to COSMIC v3 (Tate et al. 2019), were normalised to give an average exposure of each signature across the samples. The trinucleotide spectra associated with these three signatures were then mixed according to the average exposures to construct a representative POLE mutation spectrum. To construct the representative non-hypermutant CRC mutation spectrum, we used CRC samples without a pathogenic POLE mutation and, to avoid using hypermutant tumours, with a total number of single base substitutions (as given by the spectra) of less than 50,000. This threshold was determined via k-means clustering on the mutation counts with k = 2, which resulted in 44 samples. The resulting trinucleotide counts (accessible at www.synapse.org/#!Synapse:syn11726620) were then averaged to give the representative spectrum.

Assuming either non-hypermutant or POLE-mutated representative mutational spectra, the proportion of new mutations resulting in a given KRAS variant was modelled as follows. For a variant of class X[A>B]Y, the numerical value of the relevant channel of the mutational spectrum was divided by the number of genome-wide contexts of type XAY, under the rationale that the probability that a new mutation is a given variant is equal to the product of the probability that the new mutation is of a given class and the probability that it falls at that genomic location. Genome-wide trinucleotide counts were obtained by applying the function trinucleotide frequency from the R package Biostrings (Pagès et al. 2024) to the primary assembly elements of the R package BSgenome.Hsapiens.UCSC.hg38.

### Classification of *KRAS* driver variants as strong or weak

A curated list of KRAS driver variants was compiled based on multiple data sources, including intOGen (Gonzalez-Perez et al. 2013), cBioPortal (Cerami et al. 2012), COSMIC (Tate et al. 2019), the UK 100,000 Genomes Project (Cornish et al. 2024), and annotations from functional studies. The following criteria were used to classify variants as weak or strong: (i) Codon 12 and 13 variants were labelled as strong, and variants in other exons were labelled as weak (Loree et al. 2021); (ii) Bioactivation scores, as defined in (Haigis 2017) and based on biochemical data from (Hunter et al. 2015), were clustered using k-means with k = 2. Variants with high bioactivation scores were labelled as strong, while variants with low bioactivation scores or without a score were labelled as weak. Variants classified as strong under this criterion were p.G12D and p.G12V. (iii) We estimated the expected number of variants under mutational processes alone by multiplying the representative standard colorectal cancer mutation spectra by the number of KRAS variants observed in cBioPortal data from colorectal cancers without a *POLE* mutation. The ratio of observed to expected variants was then clustered using k-means, and variants with high observed-to-expected ratios were annotated as strong, while the remainder were labelled as weak. Variants classified as strong under this criterion were p.G12D, p.G12V, p.G12A, and p.G13D. Restricting the analysis to microsatellite-stable (MSS) samples resulted in only p.G12D and p.G12V being classified as strong drivers.

## Acknowledgements

We thank Ignacio Soriano, Steve Thorn, and Meritxell Brunet Guasch for helpful discussions. We gratefully acknowledge the participants of the National Genomic Research Library (NGRL), whose contributions made this research possible. Secure access to the NGRL under project ID 26 was provided by Genomics England, which delivers the NGRL in partnership with NHS England, and is wholly owned by the UK Department of Health and Social Care. The NGRL contains participants’ health data collected by the NHS as part of their care, along with samples and data from their participation in research, for which fully informed consent has been obtained. This includes genomic and clinical data provided through the NHS Genomic Medicine Service, as well as data obtained through research studies, including the 100,000 Genomes Project and the Generation Study, both of which are delivered in partnership with the NHS, and from other research cohorts involving external collaborators.

## Author contributions

M.D.N and I.T. designed the research; M.D.N performed the analysis; M.D.N and I.T. wrote the manuscript.

## Data availability

With the exception of the data from the 100,000 Genomes Project, all data analysed in this study are publicly available in the referenced repositories. For the 100,000 Genomes Project data: Data from the National Genomic Research Library (NGRL) used in this research are available within the secure Genomics England Research Environment. Access to NGRL data is restricted to adhere to consent requirements and protect participant privacy. Data used in this research include: the *KRAS* codon mutated in colorectal cancer samples, annotated by mutational subtype, derived from the Genomic Data Table of (Cornish et al. 2024). Access to NGRL data is provided to approved researchers who are members of the Genomics England Research Network, subject to institutional access agreements and research project approval under participant-led governance. For more information on data access, visit: https://www.genomicsengland.co.uk/research

## Code availability

Code for all analyses is available at https://github.com/MichaelDNicholson/hypermutantWeakDrivers.

## Supplementary Legends

Supplementary note: details of mathematical model, analytical derivations, plausible parameter range, and approximate Bayesian computation.

Supplementary figure 1: Contingency tables, odds ratios, and p-values (Fisher’s test) for different strong/weak criteria for *KRAS* driver variants in the cBioPortal data.

Supplementary figure 2: Analogous to Figure 6A-C for different criteria of classifying *KRAS* variants as weak or strong, which changes the mutational biases to the driver class. For each row we show: (left) heatmap displaying parameters such that the expected pathway to the third driver differs or remains the same between non-hypermutant and *POLE*-mutant CRCs, using estimated mutational biases to weak and strong *KRAS* drivers (Fig. 5D); (centre) as in the left plot, with equal mutation rates between subtypes; (right) as in the left plot, assuming the tertiary driver rate in non-hypermutant CRCs = 10^−5^. Title above rows is the criteria used to classify *KRAS* variants as strong drivers.

Supplementary figure 3: (left) Prior samples of the weak driver mutation rate in non-hypermutant used for the approximate Bayesian computation analysis; (right) Posterior samples of the weak driver mutation rate in non-hypermutant resulting from the approximate Bayesian computation analysis.

Supplementary table 1: *KRAS* mutation data from colorectal cancer samples, provided by the listed studies, were downloaded from the cBioPortal data portal and from the Genomics England Research Environment.

Supplementary table 2: *KRAS* driver variants considered in this study and their classifications as weak or strong drivers, based on criteria as explained in supplementary information.

