## Supplementary table 1 for "Hyper-mutational processes provide a head-start for weak cancer drivers: explaining atypical *KRAS* variants"

| Study | Samples |
|---|---|
| **cBioPortal: bowel_colitis_msk_2022** | 89 |
| **cBioPortal: coad_caseccc_2015** | 29 |
| **cBioPortal: coad_cptac_2019** | 110 |
| **cBioPortal: coadread_dfci_2016** | 619 |
| **cBioPortal: coadread_genentech** | 74 |
| **cBioPortal: coadread_mskcc** | 138 |
| **cBioPortal: coadread_mskresistance_2022** | 22 |
| **cBioPortal: coadread_tcga** | 640 |
| **cBioPortal: coadread_tcga_pan_can_atlas_2018** | 594 |
| **cBioPortal: coadread_tcga_pub** | 276 |
| **cBioPortal: crc_apc_impact_2020** | 471 |
| **cBioPortal: crc_dd_2022** | 47 |
| **cBioPortal: crc_msk_2017** | 1134 |
| **cBioPortal: crc_nigerian_2020** | 64 |
| **cBioPortal: rectal_msk_2019** | 339 |
| **cBioPortal: rectal_msk_2022** | 801 |
| **100,000 genomes project V8 (only with oncogenic KRAS)** | 816 |

Supplementary table 1: *KRAS* mutation data from colorectal cancer samples provided by the listed studies were downloaded from the cBioPortal data portal and from the Genomics England Research Environment.
