## Supplementary table 2 for "Hyper-mutational processes provide a head-start for weak cancer drivers: explaining atypical *KRAS* variants"

| mutCds | mutAA | triContext | mutType | boostDM | codon | relMutRateCrcPole | relMutRateCrc | relMutRateMSI | haigisActivationScore | classActScore | class1213 | classCbioCnt | classCbioCntMssStrict |
|---|---|---|---|---|---|---|---|---|---|---|---|---|---|
| **c.35G>A** | p.G12D | ACC | A[C>T]C | TRUE | 12 | 3.62239079403612E-11 | 1.26539687391834E-10 | 2.00873845495349E-10 | 5130 | Strong | Strong | Strong | Strong |
| **c.35G>T** | p.G12V | ACC | A[C>A]C | TRUE | 12 | 4.43596182839995E-11 | 1.69481699622471E-10 | 5.35602218848631E-11 | 1726.2 | Weak | Strong | Strong | Strong |
| **c.34G>T** | p.G12C | CCA | C[C>A]A | TRUE | 12 | 2.13062918431402E-11 | 1.71464299371741E-10 | 6.22217428590003E-11 | 3283 | Strong | Strong | Weak | Weak |
| **c.35G>C** | p.G12A | ACC | A[C>G]C | TRUE | 12 | 1.47299056343287E-12 | 4.9765478190354E-11 | 3.50648720519989E-11 | 165.1 | Weak | Strong | Strong | Weak |
| **c.34G>A** | p.G12S | CCA | C[C>T]A | TRUE | 12 | 1.18160795390179E-12 | 1.055209390569E-10 | 9.10190160060388E-11 | NA | Weak | Strong | Weak | Weak |
| **c.34G>C** | p.G12R | CCA | C[C>G]A | TRUE | 12 | 3.16113917279636E-13 | 2.85774878890389E-11 | 2.23179653509995E-11 | 626.4 | Weak | Strong | Weak | Weak |
| **c.38G>A** | p.G13D | GCC | G[C>T]C | TRUE | 13 | 1.42844955650311E-10 | 1.73930360392434E-10 | 6.37296093474457E-10 | 1238.4 | Weak | Strong | Strong | Weak |
| **c.37G>T** | p.G13C | CCA | C[C>A]A | TRUE | 13 | 2.13062918431402E-11 | 1.71464299371741E-10 | 6.22217428590003E-11 | NA | Weak | Strong | Weak | Weak |
| **c.37G>C** | p.G13R | CCA | C[C>G]A | FALSE | 13 | 3.16113917279636E-13 | 2.85774878890389E-11 | 2.23179653509995E-11 | NA | Weak | Strong | Weak | Weak |
| **c.38G>C** | p.G13A | GCC | G[C>G]C | FALSE | 13 | 1.05727839818753E-12 | 4.22082807957858E-11 | 3.38044487282723E-11 | NA | Weak | Strong | Weak | Weak |
| **c.40G>A** | p.V14I | ACG | A[C>T]G | TRUE | 14 | 3.15746280783741E-10 | 4.51795635394046E-09 | 3.97129778398875E-09 | NA | Weak | Weak | Weak | Weak |
| **c.57G>T** | p.L19F | TCA | T[C>A]A | TRUE | 19 | 4.56677959865201E-10 | 2.41119142293691E-10 | 4.63834359092911E-11 | NA | Weak | Weak | Weak | Weak |
| **c.57G>C** | p.L19F | TCA | T[C>G]A | TRUE | 19 | 5.0656627325839E-13 | 6.7650379355931E-11 | 2.63649771456482E-11 | NA | Weak | Weak | Weak | Weak |
| **c.64C>A** | p.Q22K | ACA | A[C>A]A | FALSE | 22 | 1.16251920625484E-11 | 1.93235767894909E-10 | 4.25048801663535E-11 | NA | Weak | Weak | Weak | Weak |
| **c.99T>G** | p.D33E | ATC | A[T>G]C | FALSE | 33 | 6.50099468592995E-11 | 1.7950428131891E-11 | 1.4807468546693E-11 | NA | Weak | Weak | Weak | Weak |
| **c.99T>A** | p.D33E | ATC | A[T>A]C | FALSE | 33 | 3.78235366757319E-12 | 5.54923094650923E-11 | 5.26132346504455E-11 | NA | Weak | Weak | Weak | Weak |
| **c.101C>T** | p.P34L | CCA | C[C>T]A | TRUE | 34 | 1.18160795390179E-12 | 1.055209390569E-10 | 9.10190160060388E-11 | NA | Weak | Weak | Weak | Weak |
| **c.101C>G** | p.P34R | CCA | C[C>G]A | TRUE | 34 | 3.16113917279636E-13 | 2.85774878890389E-11 | 2.23179653509995E-11 | NA | Weak | Weak | Weak | Weak |
| **c.169G>A** | p.D57N | TCG | T[C>T]G | FALSE | 57 | 7.30979868909882E-09 | 2.28019549233063E-09 | 1.61670356516155E-09 | NA | Weak | Weak | Weak | Weak |
| **c.176C>A** | p.A59E | GCA | G[C>A]A | TRUE | 59 | 9.80561990675538E-12 | 3.22514505165099E-10 | 4.62257172448083E-11 | NA | Weak | Weak | Weak | Weak |
| **c.175G>A** | p.A59T | GCT | G[C>T]T | FALSE | 59 | 1.71323141220487E-10 | 1.4447089175461E-10 | 4.15453169809831E-10 | NA | Weak | Weak | Weak | Weak |
| **c.179G>A** | p.G60D | ACC | A[C>T]C | TRUE | 60 | 3.62239079403612E-11 | 1.26539687391834E-10 | 2.00873845495349E-10 | NA | Weak | Weak | Weak | Weak |
| **c.183A>C** | p.Q61H | CTT | C[T>G]T | TRUE | 61 | 2.36329082944057E-10 | 1.65846775458783E-10 | 3.12169596928711E-11 | NA | Weak | Weak | Weak | Weak |
| **c.183A>T** | p.Q61H | CTT | C[T>A]T | TRUE | 61 | 4.34976438948716E-12 | 8.11138889786151E-11 | 1.98036751446009E-11 | NA | Weak | Weak | Weak | Weak |
| **c.182A>T** | p.Q61L | TTG | T[T>A]G | TRUE | 61 | 4.10154613547327E-12 | 2.71697103842738E-11 | 1.71392295651026E-11 | 107.2 | Weak | Weak | Weak | Weak |
| **c.182A>G** | p.Q61R | TTG | T[T>C]G | TRUE | 61 | 6.22959836643168E-12 | 4.7624775213001E-11 | 1.12951694300191E-10 | NA | Weak | Weak | Weak | Weak |
| **c.182A>C** | p.Q61P | TTG | T[T>G]G | TRUE | 61 | 3.76567362332351E-11 | 4.00128387629334E-11 | 2.88797059071623E-11 | NA | Weak | Weak | Weak | Weak |
| **c.181C>G** | p.Q61E | TCA | T[C>G]A | TRUE | 61 | 5.0656627325839E-13 | 6.7650379355931E-11 | 2.63649771456482E-11 | NA | Weak | Weak | Weak | Weak |
| **c.181C>A** | p.Q61K | TCA | T[C>A]A | TRUE | 61 | 4.56677959865201E-10 | 2.41119142293691E-10 | 4.63834359092911E-11 | NA | Weak | Weak | Weak | Weak |
| **c.351A>T** | p.K117N | ATT | A[T>A]T | TRUE | 117 | 3.10376418373657E-11 | 1.02282891845063E-10 | 8.9341534471008E-11 | NA | Weak | Weak | Weak | Weak |
| **c.350A>G** | p.K117R | TTT | T[T>C]T | FALSE | 117 | 9.85601525806229E-12 | 5.77589170440001E-11 | 1.59018983932647E-10 | NA | Weak | Weak | Weak | Weak |
| **c.351A>C** | p.K117N | ATT | A[T>G]T | FALSE | 117 | 2.35884837058572E-10 | 4.40035902794003E-11 | 2.1874263276468E-11 | NA | Weak | Weak | Weak | Weak |
| **c.436G>A** | p.A146T | GCT | G[C>T]T | TRUE | 146 | 1.71323141220487E-10 | 1.4447089175461E-10 | 4.15453169809831E-10 | NA | Weak | Weak | Weak | Weak |
| **c.437C>T** | p.A146V | GCA | G[C>T]A | TRUE | 146 | 1.10818917528406E-11 | 1.33101336610765E-10 | 5.10048895384325E-10 | NA | Weak | Weak | Weak | Weak |
| **c.436G>C** | p.A146P | GCT | G[C>G]T | FALSE | 146 | 5.26495896343364E-13 | 4.00916029735082E-11 | 3.5131634434728E-11 | NA | Weak | Weak | Weak | Weak |
| **c.439A>G** | p.K147E | TTT | T[T>C]T | FALSE | 147 | 9.85601525806229E-12 | 5.77589170440001E-11 | 1.59018983932647E-10 | NA | Weak | Weak | Weak | Weak |

Supplementary table 2: *KRAS* driver variants considered in this study and their classifications as weak or strong drivers based on criteria as explained in supplementary information.
