## Supplementary note for "Hyper-mutational processes provide a head-start for weak cancer drivers: explaining atypical *KRAS* variants"

#### Contents

**1 Model introduction** **1**

**2 Simulation** **2**

**3 Average time until tertiary driver** **2**

**4 Fastest path to tertiary driver** **4**

**5 KRAS specific simulation analysis** **4**

**6 Mutation and selection parameter range** **5**

#### 1 Model introduction

The setup of the model is as follows. At time $t = 0$, a tumour is initiated with a single precursor cell which divides at a rate $b_{\text{pre}}$. While we use the term ‘cell’, one can consider ‘cells’ to represent the fundamental evolutionary unit in early tumorigenesis for specific tissues, e.g. colonic crypts in which case $b_{\text{pre}}$ could represent the fission rate of bi-allelic *APC* mutant colonic crypts.

In this work we specify two classes of driver mutations, weak and strong drivers. At cell division a daughter cell acquires either a weak driver with probability $\mu_{\text{weak}}$, a strong driver with probability $\mu_{\text{strong}}$, or neither with probability $1 - \mu_{\text{weak}} - \mu_{\text{strong}}$. In the main text (Methods) we outline how to estimate such probabilities for a given set of weak and strong driver variants using total mutation burden expected per division, the trinucleotide sequence context of the drivers, and the mutational processes active in the tumour. Acquiring weak drivers increases cells’ division rate by a factor of $1 + s_{\text{weak}}$, while strong drivers increases by $1 + s_{\text{strong}}$.

Our model is designed to analyse when changes to mutational processes, which control $\mu_{\text{weak}}$ and $\mu_{\text{strong}}$, leads to weak drivers instead of strong drivers being observed in sequenced cancers. Hence a stopping criteria describing when cancer has arisen within a nascent tumour must be adopted. In the colorectal setting, we assume the precursor population to be initiating an adenomatous polyp. Within an adenoma which ultimately leads to a sequenced cancer, further driver mutation acquisition will occur, potentially down varied lineages. We suppose that after a critical number of driver events, not necessarily restricted to single nucleotide variants, a carcinoma initiating cell arises. In line with 'big bang' models of colorectal cancer intiation [1] we suppose the progeny of the founding carcinoma cell initiates an effective selective sweep (such that cells without the critical driver events are at such low frequencies as to undetectable in sequenced cancers).

For simplicity and to illustrate our main points, we focus on the setting where the critical number of driver mutations acquired is 3, e.g. representing, *APC* loss, a *KRAS* mutation and the loss of the second copy of *TP53*. In this setting, we refer to the mutation leading to carcinoma as the tertiary driver event, which occurs to a daughter cell per division with probability $\mu_3$. Motivated by *KRAS* we focus on whether the first driver acquired during growth is weak or strong. We assume a cell can only have either a weak or a strong driver, and that each type of driver arises only once.

Let $\tau_3^{(\text{weak})}$ denote the time until the tertiary driver event through cells with the weak driver mutation, and $\tau_3^{(\text{strong})}$ be analogous but through cells with the strong driver mutation. Then, weak drivers will be observed in sequenced cancers if $\tau_3^{(\text{weak})} < \tau_3^{(\text{strong})}$.

### 2 Simulation

Due to the computational cost of full stochastic simulations to realistic tumour sizes, we used an approximate scheme, based on the following general principle. Let $Z_t$ be the number of cells at time $t$ in a pure birth process initiated by a single cell at time $t = 0$ with birth rate $b$. If at birth events a mutation can occur with probability $\mu$, then mutations occur in the population at rate $(b\mu Z_s)_{s\geq 0}$. At large times $Z_t \approx We^{bt}$, where $W \sim \text{Expo}(1)$ [2], and so the arrival times of mutation events occur approximately at rate $(b\mu We^{bs})_{s\geq 0}$.

Based on this principle, for simulating mutation times in a group of cells (precursor, weak, or strong driver clone), we first sampled an exponential rate 1 random variable $W$, and then sampled the first arrival time for a Poisson process with rate $(b\mu We^{bs})_{s\geq 0}$, where $b$ and $\mu$ are specific to the population under consideration. When displaying clone sizes as a function of total population size, the preceding procedure was carried out resulting in mutation times, before numerically determining the time at which the total population (aggregating over relevant cell populations) reached the specified tumour size, which then immediately yielded sizes of given cell populations at that time.

### 3 Average time until tertiary driver

The time to initiate the weak driver or strong driver clone is partially dependent on the stochastic growth of the precursor population, thus correlations in time to tertiary driver through both paths exist. We account for this by considering the expected time to tertiary driver conditioned on the precursor population growth, and then seeking conditions such that the expected time is faster through the weak driver path.

We first state a known result on branching processes. Consider a pure birth process, initiated from a single cell, with divisions occurring without mutation at rate $b$ and divisions yielding a mutant and non-mutant cell occurring at rate $\nu$. Let $Z(s)$ be the number of cells in the pure birth process at time $s$. Then, conditional on $(Z(s))_{s\geq 0}$, for small $\nu$, the distribution of the time until the first mutation, $\tau$, is approximately

$$\mathbb{P}(\tau > t|(Z(s))_{s\geq 0}) \approx \exp\left(-\frac{W\nu}{b}e^{bt}\right), \tag{1}$$

where $W = \lim_{t\to\infty} e^{-bt}Z(t)$, and $W \sim \text{Expo}(1)$, see e.g. [3] page 82 or more generally Corollary 3 in [4]. Eq. (1) is a negative Gumbel distribution. Thus, using known properties

$$\mathbb{E}[\tau|(Z(s))_{s\geq 0}] \approx \frac{1}{b}\log\left(\frac{b}{\nu W}\right) - \gamma_E/b, \tag{2}$$

where $\gamma_E$ is Euler's constant. Averaging over $W$, or using Eq. 24 in [3]

$$\mathbb{E}[\tau] \approx b^{-1}\log(b/\nu). \tag{3}$$

Thus by adding Eq. (2) and Eq. (3), suitably parameterised, the expected time till the tertiary driver, conditional on the precursor growth can be approximately obtained.

In the setup of the present model, due to our focus on replicative polymerases with impaired proofreading, we specify mutations happen at division events with certain probabilities (although qualitatively similar conclusions will hold if mutation is uncoupled to division). So, precursor cells divide without mutation at rate $b_{\text{pre}}(1-\mu_{\text{weak}}-\mu_{\text{strong}})$ and divide producing a cell with a weak driver with rate $b_{\text{pre}}\mu_{\text{weak}}$, a strong driver with rate $b_{\text{pre}}\mu_{\text{strong}}$. Cells with weak or strong drivers divide without mutation at rate $b_{\text{pre}}(1+s_{\text{weak}})(1-\mu_3)$ and $b_{\text{pre}}(1+s_{\text{strong}})(1-\mu_3)$ respectively. Cells with a weak driver produce cells containing the tertiary driver at rate $b_{\text{pre}}(1+s_{\text{weak}})\mu_3$, and analogously for strong driver cells at rate $b_{\text{pre}}(1+s_{\text{strong}})\mu_3$.

Let the number of precursor cells at time $t$ be $Z_p(t)$, $\mathcal{Z}_p = (Z_p(t))_{t\geq 0}$ and $W_p = \lim_{t\to\infty} e^{-b_{\text{pre}}(1-\mu_{\text{weak}}-\mu_{\text{strong}})t}Z_p(t)$. By combining Eq. (2) and Eq. (3), the expected time until the tertiary driver event occurs via a weak driver, conditional on the precursor growth, is

$$\mathbb{E}[\tau_3^{(\text{weak})}|\mathcal{Z}_p] \approx \frac{1}{b_{\text{pre}}(1-\mu_{\text{weak}}-\mu_{\text{strong}})}\log\left(\frac{b_{\text{pre}}(1-\mu_{\text{weak}}-\mu_{\text{strong}})}{W_p b_{\text{pre}}\mu_{\text{weak}}}\right) - \frac{\gamma_E}{b_{\text{pre}}(1-\mu_{\text{weak}}-\mu_{\text{strong}})}$$
$$+ \frac{1}{b_{\text{pre}}(1+s_{\text{weak}})(1-\mu_3)}\log\left(\frac{b_{\text{pre}}(1+s_{\text{weak}})(1-\mu_3)}{b_{\text{pre}}(1+s_{\text{weak}})\mu_3}\right).$$

Similarly the conditional expected time until the tertiary driver event via a strong driver mutation is

$$\mathbb{E}[\tau_3^{(\text{strong})}|\mathcal{Z}_p] \approx \frac{1}{b_{\text{pre}}(1-\mu_{\text{weak}}-\mu_{\text{strong}})}\log\left(\frac{b_{\text{pre}}(1-\mu_{\text{weak}}-\mu_{\text{strong}})}{W_p b_{\text{pre}}\mu_{\text{strong}}}\right) - \frac{\gamma_E}{b_{\text{pre}}(1-\mu_{\text{weak}}-\mu_{\text{strong}})}$$

$$+\frac{1}{b_{\text{pre}}(1+s_{\text{strong}})(1-\mu_3)}\log\left(\frac{b_{\text{pre}}(1+s_{\text{strong}})(1-\mu_3)}{b_{\text{pre}}(1+s_{\text{strong}})\mu_3}\right).$$

### 4 Fastest path to tertiary driver

We seek parameters such that $\mathbb{E}[\tau_3^{(\text{weak})}|\mathcal{Z}_p] < \mathbb{E}[\tau_3^{(\text{strong})}|\mathcal{Z}_p]$. These are when

$$\frac{1}{b_{\text{pre}}(1-\mu_{\text{weak}}-\mu_{\text{strong}})}\log\left(\frac{1}{\mu_{\text{weak}}}\right)+$$
$$\frac{1}{b_{\text{pre}}(1+s_{\text{weak}})(1-\mu_3)}\log\left(\frac{(1-\mu_3)}{\mu_3}\right)<$$
$$\frac{1}{b_{\text{pre}}(1-\mu_{\text{weak}}-\mu_{\text{strong}})}\log\left(\frac{1}{\mu_{\text{strong}}}\right)+$$
$$\frac{1}{b_{\text{pre}}(1+s_{\text{strong}})(1-\mu_3)}\log\left(\frac{(1-\mu_3)}{\mu_3}\right).$$

Defining the scaled selection difference as $\Delta = \frac{s_{\text{strong}}-s_{\text{weak}}}{(1+s_{\text{strong}})(1+s_{\text{weak}})}$, this is equivalent to

$$-\log\left(\frac{\mu_3}{1-\mu_3}\right)\Delta < \frac{(1-\mu_3)}{(1-\mu_{\text{weak}}-\mu_{\text{strong}})}\log\left(\frac{\mu_{\text{weak}}}{\mu_{\text{strong}}}\right).$$

Under the assumption that $\mu_3 \ll 1$ and $\mu_{\text{weak}}+\mu_{\text{strong}} \ll 1$, this condition is approximately (readily shown by Taylor expansion)

$$-\Delta\log(\mu_3) < \log\left(\frac{\mu_{\text{weak}}}{\mu_{\text{strong}}}\right). \tag{4}$$

The above Eq. (4) is Equation 1 displayed in the main text.

### 5 KRAS specific simulation analysis

For the approximate Bayesian computation analysis (Fig. 6E-G), the priors were: $10^{\text{Uniform}[-8,-4]}$ for $\mu_{\text{weak}}$ in non-hypermutant CRCs; $s_{\text{weak}}$, $s_{\text{strong}}$ both drawn from Uniform(0,20) but samples retained only if $s_{\text{weak}} < s_{\text{strong}}$. 50,000 prior samples were generated. Note that once non-hypermutant $\mu_{\text{weak}}$ is sampled all other mutational parameters can be obtained by scaling with the measured mutational biases, the fold increase in mutation burden per division, and that $\mu_3 = 10^{-5}$ in non-hypermutant simulations. For a given parameter set, we simulated the same number of non-hypermutant CRCs with KRAS as observed in the cBioPortal data, and similarly for the POLE-mutant case. Of these, we then calculated the simulation proportion for which the third driver first arrived on cells with a weak versus strong driver. These proportions were compared to the observed data, and parameters samples with total distance between the observed weak driver proportion and simulated less than 0.05, were considered as approximate posterior samples, resulting in 337 samples. For the MSI posterior predictive density (Fig. 6H), we simulated using ABC accepted parameters sets, as described above. We assumed a 10 fold change in the mutation rate per division between non-hypermutant and MSI CRCs, and used a mutational bias to weak drivers in MSI CRCs of 6.73 (estimated as explained in the main text). For each parameter set, we simulated cohorts of MSI tumours of equal size to that observed aggregating over both the cBioPortal and 100,000 Genomes Project data, and calculated the simulation proportion for which the third driver first arrived on cells with a weak versus strong driver.

### 6 Mutation and selection parameter range

Here we highlight literature which motivates the parameter regimes used for simulation and analytic analysis.

Selection: Estimates for the selective advantage provided by driver mutations during tumour growth sd, in contrast to the selective advantage of growing cells compared to wild-type cells, are recently emerging. Based on colorectal adenoma measurements in Baker et al. [5], Paterson et al. [6] estimated *KRAS* G12D provides a selective advantage of sd=0.35 to increase the colon crypt fission rate. For carcinoma samples, Williams et al. [7] estimated selective coefficients of between 0.2 and 0.8 from TCGA COAD VAF samples. Using multiregion sequencing of CRCs Househam et al. [8] report increases in growth rate of up to 20 fold, which would result in sd=19. For blood cancers, a selective coefficient of 1.41 was estimated for a *KRAS* G12C variant estimated during the expansion of CLL by Lee and Bozic [9]), while for myeloproliferative neoplasm data, Johnson et al. [10] inferred secondary drivers to have selective coefficients of around 3. Thus, based on these non-exhaustive examples, a generous parameter range for selective coefficients is (0,20].

Mutation: As we are motivated by *KRAS* mutations acquired early during colorectal tumours, we focus on stem cells acquiring driver mutations between colonic crypt fission events. Colonic crypts contain on the order of 9 stem cells [11] and each colonic stem cell accumulates approximately 40 mutations/year [12, 13]. Assuming that $APC^{-/-}$ crypts undergo fission at rate 0.2/year [5], then between fission events approximately 40*9*5 = 1800 mutations should occur within a crypt. With a uniform mutation rate over bases, and assuming each of the 3 alternative bases is equally likely, the probability of any given mutation occurring is then approximately $1800/(3*6*10^9) = 10^{-7}$. If we consider on the order of 100 weak driver sites, the probability of generating one of these between fission events is $10^{-5}$ and with a *POLE* mutation $10^{-3}$.
