## Supplementary material for "Hyper-mutational processes provide a head-start for weak cancer drivers: explaining atypical *KRAS* variants": Sup Fig 1

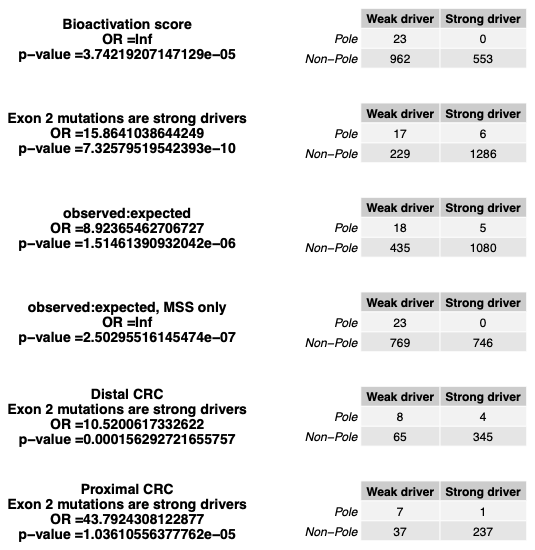

Supplementary Figure 1: Contingency tables, odds ratios, and p-values (Fisher’s test) for different strong/weak criteria for *KRAS* driver variants in the cBioPortal data.
