## Supplementary material for "Hyper-mutational processes provide a head-start for weak cancer drivers: explaining atypical *KRAS* variants": Sup FIg 2

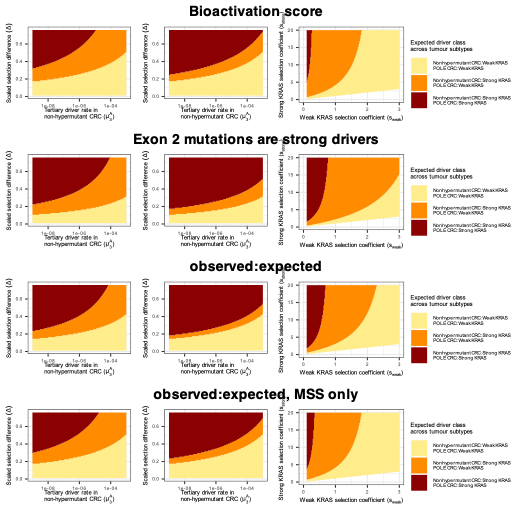

Supplementary Figure 2: Analogous to Figure 6A-C for different criteria of classifying *KRAS* variants as weak or strong, which changes the mutational biases to the driver class. For each row we show: (left) heatmap displaying parameters such that the expected pathway to the third driver differs or remains the same between non-hypermutant and *POLE*-mutant CRCs, using estimated mutational biases to weak and strong *KRAS* drivers (Fig. 5D); (centre) as in the left plot**,** with equal mutation rates between subtypes; (right) as in the left plot**,** assuming the tertiary driver rate in non-hypermutant CRCs = 10^-5^. Title above rows is the criteria used to classify *KRAS* variants as strong drivers.
