## Supplementary material for "Hyper-mutational processes provide a head-start for weak cancer drivers: explaining atypical *KRAS* variants": Sup Fig 3

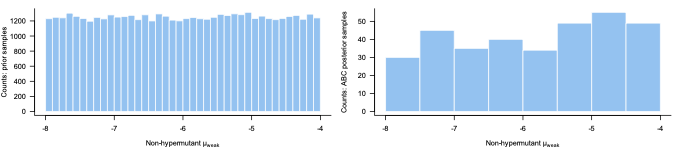

Supplementary Figure 3: (left) Prior samples of the weak driver mutation rate in non-hypermutant used for the approximate Bayesian computation analysis; (right) Posterior samples of the weak driver mutation rate in non-hypermutant resulting from the approximate Bayesian computation analysis.
